# Invertebrate species distributions in urban ecosystems are driven by habitat availability, not by anthropogenic drivers

**DOI:** 10.1101/2024.08.26.609715

**Authors:** Joeri Morpurgo, Roy P. Remme, Quinten Mudde, Emilie A. Didaskalou, Kornelia J. Serwatowska, Krijn Trimbos, Mingming Hu, Peter M. Van Bodegom

## Abstract

Rapid urbanisation puts pressure on biodiversity by habitat fragmentation and habitat quality among others. These habitat changes are known to affect species dispersal and connectivity and ultimately species distribution in many ecosystems. However, little is known about the changing urban species distributions which in turn influence community assembly. New DNA-based sampling methods combined with species distribution modelling provide a way to quantitatively estimate urban community assembly through assessing many species distributions and their drivers simultaneously.

We investigated species distributions of entire communities in the city of The Hague (the Netherlands) by sampling DNA with two distinct methods: collecting data on invertebrate occurrence with traditional trapping (bulk; n = 205) and a novel Environmental DNA method (eDNA; n = 207). After DNA sequencing, species were identified using Operational Taxonomic Units. Subsequently, individual species presence and absence were used in Species Distribution Models (SDMs), based on spatial information on vegetation and anthropogenic influences.

The results show a difference in coverage of sampling methods (bulk vs. eDNA), indicating their complementary information. The models on species distributions were generally significantly better than random models (59.5%), and performed well during calibration (90.4%, AUC > 0.70). In contrast, during validation very few SDMs (1.3%, AUC > 0.70) performed adequately in predicting species distributions.

Through this novel combination of DNA sampling with SDMs we show that density and structure of vegetation, as well as distance to water are more important for urban invertebrate distribution than direct anthropogenic pressures. This suggests that dispersal is not a limiting factor in The Hague’s urban environment. The availability of a variety of urban green infrastructures seems sufficient to attract many of the species observed. Hence, ensuring sufficient green infrastructure in the urban environment should be the first priority to enhance biodiversity in the urban environment.

**Highlights:**

- Vegetation and water indicators are the most important predictors for accurate assessment of species’ distribution
- Anthropogenic pressures have lower impact on urban species distributions
- DNA-SDM combinations show potential in guiding urban planners in evidence-based decision making

## 1. Introduction

Cities have long been considered ecological deserts. With urbanization rapidly increasing in many parts of the world, these ecological deserts increasingly replace the original habitat and local species are lost where cities are built (Güneralp and Seto, 2013). This loss in biodiversity is exacerbated by the tendency to build cities in biodiversity hotspots, with models suggesting that a third of the land-dwelling vertebrates are impacted by urbanization (Simkin et al., 2022). Within cities, new habitat is built, with usually very different properties than the original natural habitat. Compared to the natural habitat, green infrastructure (i.e. a network of natural area managed to deliver ecosystem services and protect biodiversity (European Commission, 2014)) in the urban environment exhibits uniquely high levels of anthropogenic stress in the form of management, human disturbance, exotic species, climatic extremes, and nutrient loading, among others (Peters et al., 2015; Orr et al., 2022; Vilmi et al., 2017). Urban dwelling organisms will necessarily have to deal with these abundant stressors (McCloy et al., 2024).

Despite these prevailing stressors, cities are also increasingly being recognized as refuges and potential hotspots for biodiversity, known to harbour even rare, endangered and red-listed species (Boakes et al., 2023). Simultaneously, evidence is starting to mount that core ecological factors, such as increasing habitat availability, increasing connectivity among green patches and reducing stressors, are readily enhancing urban biodiversity (Beninde et al., 2015; LaPointe et al., 2015). Several reviews and meta-analyses show that both the green infrastructure and man-made built infrastructure shape the urban biodiversity: In a meta-analysis investigating the effect of green infrastructure on urban biodiversity, covering 87 studies, species richness was best predicted by habitat richness, vegetation structure and density of green infrastructure (Beninde et al., 2015). In parallel, a review on the spatial connectivity of urban green infrastructure at city scale has been shown to be important in understanding species distributions (LaPointe et al., 2015). Subsequently, the design of grey infrastructure and anthropogenic pressures, such as urban form, policy and the socioeconomic status of neighbourhood communities also seem to be shaping urban biodiversity (Kuras et al., 2020; Schell et al., 2020). For example, small urban parks within denser urban environments have less diverse communities due to a reduction in the number of endemic and non-urban species (Amaya-espinel et al., 2019). Likewise, neighbourhoods with higher socioeconomic status tend to have higher plant and animal diversity (Leong et al., 2019). These studies indicate that urban biodiversity is moulded by cities, their green and grey infrastructures, layout and use.

Currently, the knowledge on drivers important for urban biodiversity is developing, yet empirical evidence on the drivers of individual distributions of species and their assembly into communities is mostly lacking (Pickett et al., 2011; Andrade et al., 2021). Understanding urban species environmental requirements through the design of the quantity, spatial lay-out and quality of urban green and built infrastructure is key to enhancing its biodiversity. These spatial features and their design are critical for a plethora of ecological processes such as dispersal and functioning of trophic networks. Viewing these spatial green and grey features as environmental filters can help us understand how urban species are pooled into communities through linking the regional climate, spatial and socio-economic development of the city and ecological processes in relationship to species and their functional traits (Aronson et al., 2016; Zhao et al., 2023). If traditional ecological theories developed in natural habitat apply, then conventional actions to preserve target species and their associated biodiversity should prove effective in the cities. Yet, if species and their assembly into communities differ in urban environments, we need to understand how to support urban biodiversity differently compared to natural areas. By understanding the critical features of urban community assembly, we can further elucidate the underlying mechanisms of urban biodiversity, and in turn develop guidelines that support, protect, and conserve urban target species and biodiversity more effectively (IUCN et al., 2022), ultimately finetuningthe critical design factors of urban green infrastructure to support urban biodiversity.

Invertebrates are a particularly important part of ecosystems. They are essential for ecosystem functioning and health through pollination, nutrient cycling, soil bioturbation, and serving as crucial links in trophic networks (Buchholz & Egerer, 2020; Belovsky and Slade, 2000; Drager et al., 2016; Eggleton, 2020). For example, flower-visiting insects are indispensable for pollinating and sustain plant and invertebrate populations throughout the urban and peri-urban environment (Eggleton, 2020). The trophic cascade from plants to invertebrates to soil nutrients is highly important for plant diversity, soil health, and overall ecosystem health (Belovsky and Slade, 2000). Despite their evident importance, invertebrate numbers in protected areas continue to dwindle at alarming rates (Hallmann et al., 2017). This decline threatens the stability of many critical ecosystem functions (Wagner et al., 2021). Here, urban environments may also serve as genetic reservoirs as some animal species have been recorded to disperse from rural to urban environments and vice versa (Brashear et al., 2015; Varner et al., 2014). The evidence of urban environments as genetic reservoirs for invertebrates is lacking but likely, with research suggesting that urban invertebrate gene flow is seems to be species-dependent (Collins et al., 2024). Simultaneously, urban environments are known to harbour rare and endangered species different from local communities, yet their drivers are poorly known (Andrade et al., 2021; Boakes et al., 2023). Considering the above, we urgently need to understand the drivers of invertebrate species fundamental niche and their resulting distributions so that we can design cities and their layout with green infrastructure that meaningfully supports invertebrate communities.

Understanding the drivers of urban invertebrate species fundamental niches can be achieved through Species Distribution Modelling (SDM) of the species realised niches. These models estimate the importance of environmental features in the ecological landscape by modelling empirical data on species presence and absence against drivers considered to be relevant (Dutta et al., 2022; Naimi and Araújo, 2016). Such an approach is rare for entire ecosystems at the city scale, but has been shown to be successful in modelling plants, leafhoppers and grasshoppers and their driving factors (Kattwinkel et al., 2009). In parallel, recent developments in DNA techniques, specifically environmental DNA (eDNA) and bulk metabarcoding, now allow for quick sampling and standardized analysis of whole communities (Cristescu and Herbert, 2018, Gleason et al., 2021). Moreover, these DNA techniques generate presence/absence data that can subsequently be used by SDMs to assess the community assembly of an entire community while reducing sampling time significantly.

Through the combination of DNA-based data and SDMs, we aim to understand species, their distribution, and their drivers, in the urban context resulting in unprecedented coverage and scale. More specifically, we use soil eDNA and invertebrate bulk metabarcoding with SDMs to investigate the spatial distribution of invertebrates using predictors traditionally considered important for urban biodiversity. With this DNA-SDM combination, we include a broad range of urban invertebrate species and develop well-performing SDMs. Our approach enables investigating the effects of urban green infrastructure on species and their distribution and draw inferences on ecological connectivity and biodiversity hotspots. This knowledge can be translated into guidelines making it easier for urban planners and municipalities to develop strategies that effectively support and enhance urban biodiversity through target species (Kirk et al., 2021; Pierce et al., 2024).

## 2. Methods

We investigated urban invertebrate species distributions in a three-step process. First, we selected relevant sampling sites for invertebrate trapping and soil collection through conditional Latin Hypercube Sampling (cLHS). Second, we extracted, amplified, and sequenced the DNA from both bulk invertebrate samples and soil samples. Sequenced samples were processed with the DADA2 bioinformatics pipeline, clustered into OTUs and BLASTed against the BOLD reference database to assign taxonomy. Finally, we modelled the identified species and their distributions with predictors thought to be ecologically important, at multiple scales of aggregation. The resulting SDMs were used to infer ecological hotspots at the species level and assess the most important drivers of their distributions.

### 2.1 Sampling and location

Invertebrates and soil were sampled in a broad range of green infrastructure types in The Hague, The Netherlands (52_◦_04’57.0”N 4_◦_17’40.3”E). We excluded Leidschenveen in our sampling design, as it was both isolated from the most of The Hague and also logistically infeasible to get there given our sampling equipment and design. We sampled 207 sites on non-rainy days from the 15th of June until the 4th of August 2022 (Fig. 1). Sampling consisted of collecting soil and setting traps on Monday or Tuesday, and retrieving traps two days later on Wednesday and Thursday.

**Figure 1.**
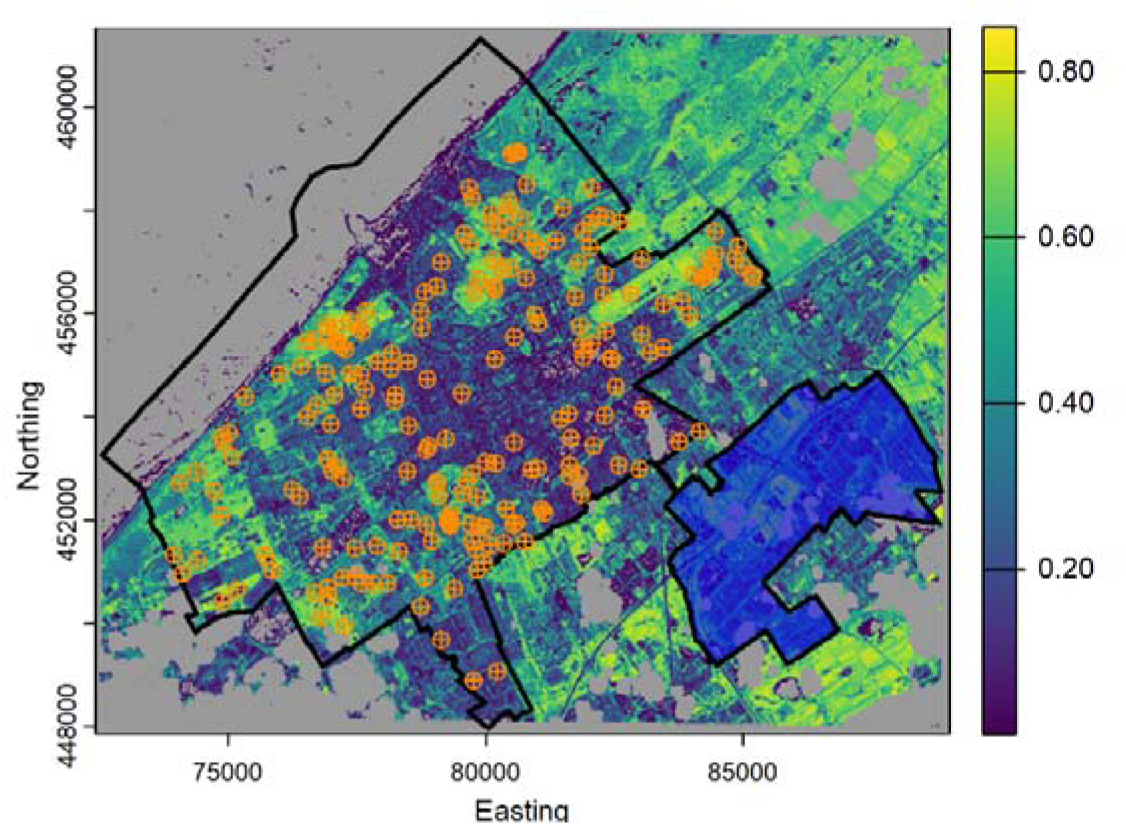
Sampling locations against a background of scaled NDVI in The Hague, The Netherlands. The orange crossed circles indicate a sample being taken there. The blue polygon indicates the excluded zone (Leidschenveen).

Sample site selection aimed to cover the full range of indicators expected to be of influence on invertebrate diversity. To ensure the full coverage of the relevant indicators, we used conditional Latin Hypercube Sampling (cLHS). This method is recommended in cases where sampling is logistically limited, as it allows for maximum coverage in feature space with the least amount of samples. The indicators were largely based on a review covering green infrastructure and its relation to several biodiversity taxa (Morpurgo et al., 2023). Our included indicators cover the basic needs for invertebrates, such as food, shelter and water. In addition, we added data on buildings and land uses to fully represent the urban landscape (Table S1).

We sampled both pitfall and pan traps and soil eDNA, to both compare methods and capture most invertebrate biodiversity through off-setting any potential trapping biases (Montgomery et al., 2021). The combination of pan and pitfall traps allows for capturing both species that have high dispersal capacity (i.e. flying insects) and species with low dispersal capacity (i.e. walking or crawling insects), which may face different dispersal barriers in the urban environment. For the soil eDNA sample, we dug up approximately 500ml of topsoil that was mixed in a plastic bag. From this mixture, a sample of 50ml soil was stored in a falcon tube and stored in a freezer at -20°C. For the invertebrate bulk sampling, one pitfall and three colours of pan traps (white, yellow, blue) were set with 0.1% perfume-free and colourless soap (Everingham, 2023). The pitfall trap was transparent and measured 80mm length by 115mm diameter, which is considered optimal given the relatively smaller size of our targeted invertebrate species (Brown and Matthews, 2016). Yellow and blue pan traps were 40mm deep and 105mm in diameter. The white pan traps were 55mm deep and 110mm in diameter. The selected colours are known to attract different pollinator species, and thus we combined them to off-set any species-colour related biases (Vrdoljak and Samways, 2012). After 48 hours, trapped invertebrates were filtered through coffee filters, and scraped into 50ml falcon tubes containing 96% ethanol and stored in a freezer at -20°C.

### 2.2 DNA extractions

To extract eDNA from soil, we added 15 ml of saturated phosphate buffer (Na_2_HPO_4_; 0.12m) to 15 grams of soil and homogenized through vigorous shaking, followed by stirring at 600 rpm for 10 minutes to further homogenize the samples (Taberlet et al., 2012). After shaking, 2 ml of the soil-buffer mixture was transferred in a 2.5ml tube and samples were further extracted using the Qiagen DNeasy PowerSoil Pro Kit following the manufacturer’s protocol starting from step 3. The resulting eDNA extracts were stored at -20℃ until further use.

We used and adjusted protocol to extract DNA from the invertebrate bulk from Beentjes et al. (2019) in the following aspects. First, we homogenized the bulk using the IKA Ultra-Turrax Tube Drive with 15 steel beads of 5mm diameter per 50 ml tube for 5 minutes at max speed. For each sample, we twice transferred 2ml of the mixture to 2.5ml tubes, each tube serving as subsample. We centrifuged the subsamples at 16000 g for 3 minutes separating the homogenized invertebrate bulk from the ethanol. After, we removed the supernatant ethanol and dried the samples at 50℃ until visually dry. The dried subsamples were combined and digested using 20_μ_L proteinase K and 180_μ_L ATL buffer for 24 hours at 56℃ and 600rpm. The digested samples were centrifuged at 10000g for 3 minutes to separate the non-digestible material from the eluted DNA. If eluted DNA did not separate, the sample was centrifuged at 16000 g for 3 min. Finally, the recovered supernatant containing the DNA was transferred onto a new tube and followed the manufacture’s protocol from step 3 onwards. Resulting DNA extracts were stored at -20℃ until further use.

### 2.3 PCR amplification and DNA sequencing

For both soil and invertebrate samples, we targeted the mitochondrial gene Cytochrome c oxidase I (COI), but used two different primer sets that previous research suggested to be optimal for assessing general invertebrate communities for soil eDNA and bulk DNA extracts (Leray et al., 2013; Elbrecht et al., 2019). For the soil eDNA extracts, we used the primer mlCOIintF and dgHCO2198 resulting in an amplicon of 313 base pairs (Table S2; Leray et al., 2013). For invertebrate bulk DNA extracts, we used the forward primer BF1 and the reverse primer BR2 resulting in an amplicon of 316 base pairs (Table S2; Elbrecht et al., 2019). Both primer sets included Illumina tails, to attach indexing sequences required for DNA sequence demultiplexing.

For the first PCR round, samples were amplified in triplicate in 25_μ_L reactions containing 2_μ_L of DNA extract, 8_μ_L of DNase/RNase free water, 12.5_μ_L Taqman, Environmental Master Mix 2.0 and 2.5_μ_L of each primer (10uM). PCR was performed in a Bio-Rad C1000 touch Thermal Cycler, using the following program: initial denaturation at 98℃ for 10 minutes, 30 cycles of denaturation at 95℃ for 15 seconds, annealing at 45℃ for 30 seconds, and extension at 72℃ for 40 seconds, followed by a final elongation at 72℃ for 5 minutes. Samples were visually checked on 2% precast agarose E-gels™ with SYBR™ Safe (Invitrogen) to ensure successful amplification before proceeding. All samples’ triplicates were combined into one library and cleaned with NucleoMag NGS Clean-up and Size Select beads (MACHEREY-NAGEL) at a ratio of 0.9:1 (beads/sample). For the second PCR round, libraries were dual indexed in 20_μ_l reactions containing 2_μ_l of library (1^st^ PCR product), 6_μ_l of DNase/Rnase free water, 10_μ_l Taqman, Environmental Master Mix 2.0 and 1.5_μ_l of each IDT10 index (10_μ_M). The PCR program used was: initial denaturation at 98℃ for 10 minutes, 8 cycles of denaturation at 95℃ for 15 seconds, annealing at 55℃ for 30 seconds, and extension at 72℃ for 40 seconds, followed by a final elongation at 72℃ for 5 minutes.

As sequencing is a random process, equimolar pooling was chosen to equalize the chances of the DNA being sequenced (Muller et al., 2019). We equimolarly pooled the library based on DNA concentrations measured with an 4200 TapeStation from Agilent using the D1000 ScreenTapes. In particular, this type of pooling dilutes samples with abundant PCR products to allow samples with little PCR product to have an equal sequencing chance. The soil and invertebrate bulk pooled libraries were then cleaned with NucleoMag NGS Clean-up and Size Select beads (MACHEREY-NAGEL) at a ratio of 0.9:1 (beads/sample). Finally, the two pooled libraries (soil & invertebrate bulk) were equimolarly pooled and sequenced by the Illumina NovaSeq 6000 SP system on a 250PE flow cell.

### 2.4 Bioinformatics

To identify the sequenced COI DNA, we used a software combination of DADA2, CutAdapt, VSEARCH, BLAST and the public BOLD reference database. First, we started with DADA2 to filter reads with a higher-than-expected error of 1, excluding low-quality reads (Callahan et al., 2016; Elbrecht et al., 2019). The primers from the filtered reads were removed using Cutadapt, and reads shorter than 300 bp were omitted as they are substantially shorter than the target amplification area (Martin, 2011). After, we used the Big Data Paired-end DADA2 workflow to generate chimera-free amplicon sequence variants, but decreasing the maxEE argument to a stricter 1. Using VSEARCH, the chimera-free amplicon sequence variants were clustered into Operational Taxonomic Units (OTUs) using the 97% similarity cut-off, typically representing a species (Elbrecht et al., 2018; Rognes et al., 2016). These OTUs were identified using BLAST with the BOLD COI public database (BOLDSYSTEMS, 2023; Altschul et al., 1990). The BLAST was set to give a maximum of 10 matches, which also had to align with the expected location of the amplicon within the COI marker. For every OTU, we filtered to a single taxonomic annotation per OTU based on sequentially the highest bitscore, lowest e-value and highest percentage similarity (>70%).

For the SDMs, we also omitted any OTU that was identified as a non-invertebrate species. To assess the impact of dispersal capacity on the SDM performance, we grouped OTUs in three groups based on estimated dispersal capacity by taxonomic groups. The dispersal groups were invertebrates that primarily fly (flying; *Diptera* and *Hymenoptera*), facultative flying invertebrates that fly and crawl (facultative flying; *Coleoptera, Dermaptera, Hemiptera* and *Orthoptera*) and ground-dwelling invertebrates (ground; *Annelida, Arachnida, Collembola, Diplopoda, Isopoda* and *Symphyla*).

### 2.5 Species distribution models

We used SDMs based on the presence or absence of the identified species from either the eDNA or bulk metaBarcoding datasets to infer invertebrate species distributions. For robust SDMs, we chose a minimum of 15 presences (out of the 200+ samples) for an OTU to be modelled. The SDMs were modelled across eight scales (10m^2^, 20m^2^, 30m^2^, 40m^2^, 50m^2^, 100m^2^, 200m^2^ and 500m^2^) and two sets of drivers. The first set of drivers focused on land use variables as identified in CUGIC (Consolidated Urban Green Infrastructure Classification, Morpurgo et al. 2023):

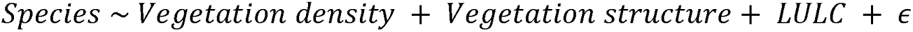

The second set of drivers, hereafter called ECO, includes more predictors that are considered ecologically important to invertebrates:

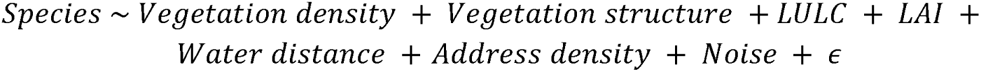

Where in both formulas *Species* denotes the presence or absence of an identified invertebrate species. The term *Vegetation density* contains numerical information between 0 *—* 1 on the density of foliage in a given cell, using scaled NDVI (Carlson and Ripley, 1997). The term *Vegetation structure* is a numerical indicator between 1 and 2, where 1 describes all vegetation being below 1 meter high, and 2 describes all vegetation being above 1 meter high. Upon aggregation to a larger spatial scale, this term describes the proportion of low vs. high vegetation in a cell. The land use and land cover is described categorically in the *LULC* term, including I) natural, II) park, III) residential and IV) buffer zone. For the ECO model, we also included numerical information on leaf area index by the term *LAI*, and distance to water by the term *Water Distance* to include data on species’ needs (Pfeifer et al., 2012; Crocker et al., 2017). *Address density* and *Noise* are indicators reflecting the level of anthropogenic influence that poses potential barriers to species (Den Dulk et al., 1992; Van Der Waals, 2002). The final E term encompasses the remaining error in both the CUGIC and ECO models.

The SDMs used 70% of the data to calibrate the model and used 30% of that to validate the accuracy of the model. All SDMs were run in the R package SDM, using several model structures (GAM, MDA, FDA, MARS, CART, MAXENT) and replicated 10 times to increase the robustness of the analysis (Naimi and Araújo, 2016). This resulted in 960 models trained and validated per OTU, including 2 sets of drivers, 6 model structures, 8 spatial scales and 10 replications. Accuracy was evaluated using traditional AUC values thresholds against the mean of the 480 validation AUCs per OTU (Gorunescu, 2011).

### 2.6 Drivers of SDM performance

To evaluate the drivers of invertebrate species we investigated the drivers of the validation AUCs. For every SDM, the validation AUC was compared to 100 models with identical settings, but using random presence/absence data (i.e., a random model) to assess if the SDM performed better than 95% confidence interval of the random models (Raes and ter Steege, 2007). All AUCs of SDMs that were better than random were used in a linear mixed model to assess if scale, dispersal group (flying, facultative flying, ground) or the set of drivers (CUGIC or ECO) were important drivers of AUC. For dispersal, we added an interaction with model type and scale as the drivers or spatial scale may have different effects on species with low dispersal capacity compared to species with high dispersal capacity (Wunderlich et al., 2022). Additionally, this model accounts for the effect of the calibration AUC and controls for species average differences by the random effects. The resulting model was analysed with an Tukey post-hoc test to assess difference between dispersal groups and model types. This leads to the following model:

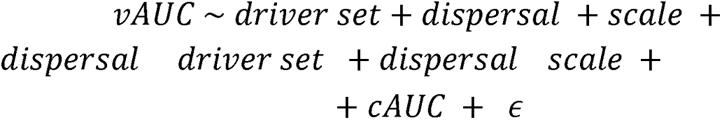

Where *vAUC* describes the mean validation AUC from the SDMs for a particular species, and is predicted by the rest of the terms. The term describes the random effects for every species, cancelling out average differences in AUC between species. The term *cAUC* contains the information of the AUC during the calibration phase.

### 2.7 Variable importance of SDM performance

To assess general variable importance we extracted the AUC variable importance from every SDM. This importance indicates how much the AUC of a model is affected by the exclusion of the excluded variable (Naimi and Araújo, 2016). For every set of drivers and dispersal groups, we assessed the differences between variable importance using an ANOVA followed by a Tukey post-hoc test. Finally, general variable importance was calculated by the frequency of a variable being chosen to be the most impactful across driver sets and dispersal groups.

**Table 1.** Description of the variables used in the Species Distribution Models and their method of aggregation. The variables LAI, Water distance, Address density and noise only apply to the ECO model. Aggregation methods describe either a mean (Bilinear), a sum method or the most frequent value (modal). Data statistics are identical for both the soil and bulk dataset, unless stated differently.

| Variable at 10m<sup>2</sup> | Description | Aggregation method | 20m<sup>2</sup> | 30m<sup>2</sup> | 40m<sup>2</sup> | 50m<sup>2</sup> | 100m<sup>2</sup> | 200m<sup>2</sup> | 500m<sup>2</sup> |
| --- | --- | --- | --- | --- | --- | --- | --- | --- | --- |
| Species (Reponse variable) | A binary variable representing the presence (1) or absence (0) of an OTU identified to a species at >97% identical DNA-match with the BOLD public data package (BOLDSYSTEMS, 2023). The soil eDNA contained 47 OTUs, and the invertebrate bulk contained 163 OTUs. | No aggregation | No aggregation | No aggregation | No aggregation | No aggregation | No aggregation | No aggregation | No aggregation |
| Vegetation density | A numerical variable that uses the average normalized difference vegetation index (NDVI), retrieved from Sentinel-2 on June — September 2022, and scaled accordingly to both the NDVI values found in our sample sites (NDVI: Min = 0.05; Max = 0.90). The range of scaled vegetation densities was 0.01 — 0.75, with an average value of 0.36 and a SD of 0.20. | Bilinear | Min=0.03<br>Avg=0.37<br>SD=0.20<br>Max=0.77 | Min=0.01<br>Avg=0.36<br>SD=0.20<br>Max=0.77 | Min=0.03<br>Avg=0.37<br>SD=0.19<br>Max=0.76 | Min=0.03<br>Avg=0.36<br>SD=0.19<br>Max=0.75 | Min=0.03<br>Avg=0.35<br>SD=0.17<br>Max=0.75 | Min=0.08<br>Avg=0.36<br>SD=0.16<br>Max=0.74 | Min=0.09<br>Avg=0.35<br>SD=0.14<br>Max=0.63 |
| Vegetation height | A binary variable that uses Vegetation density in combination with AHN4 LiDAR data to classify vegetation below 1m (1) or vegetation with over 10% above 1m (2) (AHN, 2024). Upon aggregation, cells were averaged to create an indicator on vegetation structure, between 0 and 1. The minimum (indicating all vegetation <1m) and maximum values (all vegetation above >1m) were 1 and 2, respectively. The average value was 1.4 and SD 0.49. | Bilinear | Min=1<br>Avg=1.45<br>SD=0.41<br>Max=2 | Min=1<br>Avg=1.45<br>SD=0.37<br>Max=2 | Min=1<br>Avg=1.46<br>SD=0.34<br>Max=2 | Min=1<br>Avg=1.44<br>SD=0.31<br>Max=2 | Min=1<br>Avg=1.45<br>SD=0.27<br>Max=2 | Min=1<br>Avg=1.45<br>SD=0.25<br>Max=1.99 | Min=1<br>Avg=1.45<br>SD=0.20<br>Max=1.83 |
| LULC (land-use/landcover) | A categorical variable with four classes that describe the respective classes in the CUGIC classification (Morpurgo et al., 2023). The classes are Park (SOIL=51; BULK=50), Natural (Nat, SOIL=27; BULK=26), Residential (Resi, SOIL=101; BULK=100) and Buffer zone (Buff, SOIL=28; BULK=29). | Modal | Park=50<br>Nat=28<br>Resi=100<br>Buff=29 | Park=51<br>Nat=28<br>Resi=100<br>Buff=28 | Park=49<br>Nat=27<br>Resi=102<br>Buff=29 | Park=51<br>Nat=27<br>Resi=103<br>Buff=26 | Park=48<br>Nat=25<br>Resi=107<br>Buff=27 | Park=42<br>Nat=27<br>Resi=112<br>Buff=26 | Park=46<br>Nat=34<br>Resi=104<br>Buff=23 |
| LAI (Leaf Area Index) | A numerical variable describing the Leaf Area Index as a supplementary vegetation indicator retrieved from SNAP with Sentinel-2 imagery (European Space Agency (ESA), 2023). The range of this value is 0.02 — 2.95 with a mean of 0.51 and a SD of 0.37. | Bilinear | Min=0.07<br>Avg=0.54<br>SD=0.37<br>Max=2.85 | Min=0.02<br>Avg=0.52<br>SD=0.34<br>Max=2.15 | Min=0.10<br>Avg=0.55<br>SD=0.32<br>Max=2.32 | Min=0.08<br>Avg=0.53<br>SD=0.30<br>Max=2.25 | Min=0.07<br>Avg=0.55<br>SD=0.30<br>Max=1.87 | Min=0.11<br>Avg=0.55<br>SD=0.27<br>Max=1.67 | Min=0.15<br>Avg=0.52<br>SD=0.24<br>Max=1.66 |
| Water distance | A numerical variable describing the distance to the nearest fresh-water (Kadaster, 2022b). The range of the values are 0 — 782.6. The SOIL mean was 147.7 with a SD of 157.8. The BULK mean was 148.7 with a SD of 157.9 | Bilinear | Min=1<br>Avg=147<br>SD=157<br>Max=781 | Min=0<br>Avg=147<br>SD=158<br>Max=774 | Min=0<br>Avg=146<br>SD=158<br>Max=783 | Min=2<br>Avg=146<br>SD=155<br>Max=768 | Min=10<br>Avg=146<br>SD=156<br>Max=773 | Min=10<br>Avg=143<br>SD=151<br>Max=728 | Min=13<br>Avg=140<br>SD=143<br>Max=834 |
| Address density | A numerical variable describing the number of addresses found within the cell (Kadaster, 2022a). The range of these values is 0 — 8. The SOIL mean was 0.44 and SD was 1.34. For the BULK, the mean was 0.41 and the SD was 1.28. | Sum | Min=0<br>Avg=2<br>SD=3<br>Max=19 | Min=0<br>Avg=3<br>SD=5<br>Max=24 | Min=0<br>Avg=6<br>SD=9<br>Max=79 | Min=0<br>Avg=9<br>SD=12<br>Max=55 | Min=0<br>Avg=36<br>SD=42<br>Max=222 | Min=0<br>Avg=142<br>SD=136<br>Max=558 | Min=0<br>Avg=<br>SD=<br>Max= |
| Noise | A numerical variable describing the modeled amount of noise (Rijksinstituut voor Volksgezondheid en Milieu (RIVM), 2020). The range of these values was 39 — 71, with a mean of 53.7 and a standard deviation of 7.0. | Bilinear | Min=40<br>Avg=54<br>SD=7<br>Max=73 | Min=39<br>Avg=54<br>SD=7<br>Max=72 | Min=39<br>Avg=54<br>SD=7<br>Max=74 | Min=39<br>Avg=54<br>SD=7<br>Max=72 | Min=40<br>Avg=54<br>SD=7<br>Max=71 | Min=40<br>Avg=54<br>SD=6<br>Max=71 | Min=39<br>Avg=55<br>SD=5<br>Max=73 |

## 3 Results

### 3.1 Comparing sampling methods

In our samples, we found 1108 OTUs in the soil eDNA and 330 OTUs in the invertebrate bulk that occurred at >15 sites, making them suitable for the SDMs. Among those, 81 soil OTUs and 200 bulk OTUs could be annotated to an invertebrate organism above 70% similarity. For the annotated bulk OTUs, we found that most could be annotated with the reference data to the taxonomic level of species (n = 135) or order (n = 51). For the soil eDNA annotated OTUs, we found that most were identified to the taxonomic level of order (n = 42), family (n = 16), or species (n = 14). *Diptera*, *Isopoda* and *Coleoptera* were the most frequently identified taxa in the bulk invertebrate method, while *Collembola*, *Coleoptera* and *Hymenoptera* were proportionally more found in soil eDNA. Nonetheless, invertebrate bulk retrieved more OTUs for every taxa, except for *Hymenoptera*, *Annelida* and *Symphyla*. Running the analysis on a combined dataset resulted in 257 OTUs, indicating that only 24 OTUs overlapped both datasets. Especially the taxa of *Coleoptera* (+15, 63%), *Collembola* (+12, 55%) and *Hymenoptera* (+8, 50%) showed a substantial increase in coverage by combining the methods.

**Table 2.** The 10 most frequently found and identified OTUs of soil eDNA, invertebrate bulk metabarcoding and the combined dataset. Taxa described are OTUs identified at >70% match using BLAST with the BOLD database. Remaining OTUs are summed into the “Other” group.

| Taxa | Soil eDNA<br>(n = 81) | Invertebrate bulk<br>(n = 200) | Combined<br>(n = 257) |
| --- | --- | --- | --- |
| Diptera | 9 | 71 | 74 |
| Coleoptera | 17 | 24 | 39 |
| Collembola | 19 | 21 | 33 |
| Isopoda | 1 | 28 | 27 |
| Hymenoptera | 16 | 12 | 24 |
| Arachnida | 5 | 12 | 17 |
| Hemiptera | 0 | 15 | 15 |
| Annelida | 5 | 1 | 6 |
| Diplopoda | 1 | 2 | 2 |
| Symphyla | 2 | 0 | 2 |
| Other | 6 | 14 | 18 |

### 3.2 SDM predictive performance and maps

Calibrating all models (n = 4496) resulted in a high proportion of AUC scores indicating good model performance (n = 4051, 90.4%; AUC > 0.70) across driver sets. However, during model validation the model performance dropped very strongly (with only n = 57, 1.3%, having AUC > 0.70). Even so, most models did perform better than random models (n = 2664, 59.5%, p < 0.05). The SDM performance seems independent of the scale the predictors were aggregated to, as there was neither visual clustering of models with the same scales nor a statistical significant effect (Table S2). These results indicate that for both sets of drivers, the predictors likely overfit the species niches for both model types, creating a poor predictive model. This particular problem is more apparent in the ECO model, where the additional predictors seem to push the calibration AUCs higher, while the validation AUCs seem largely unaffected (Fig. 2).

**Figure 2.**
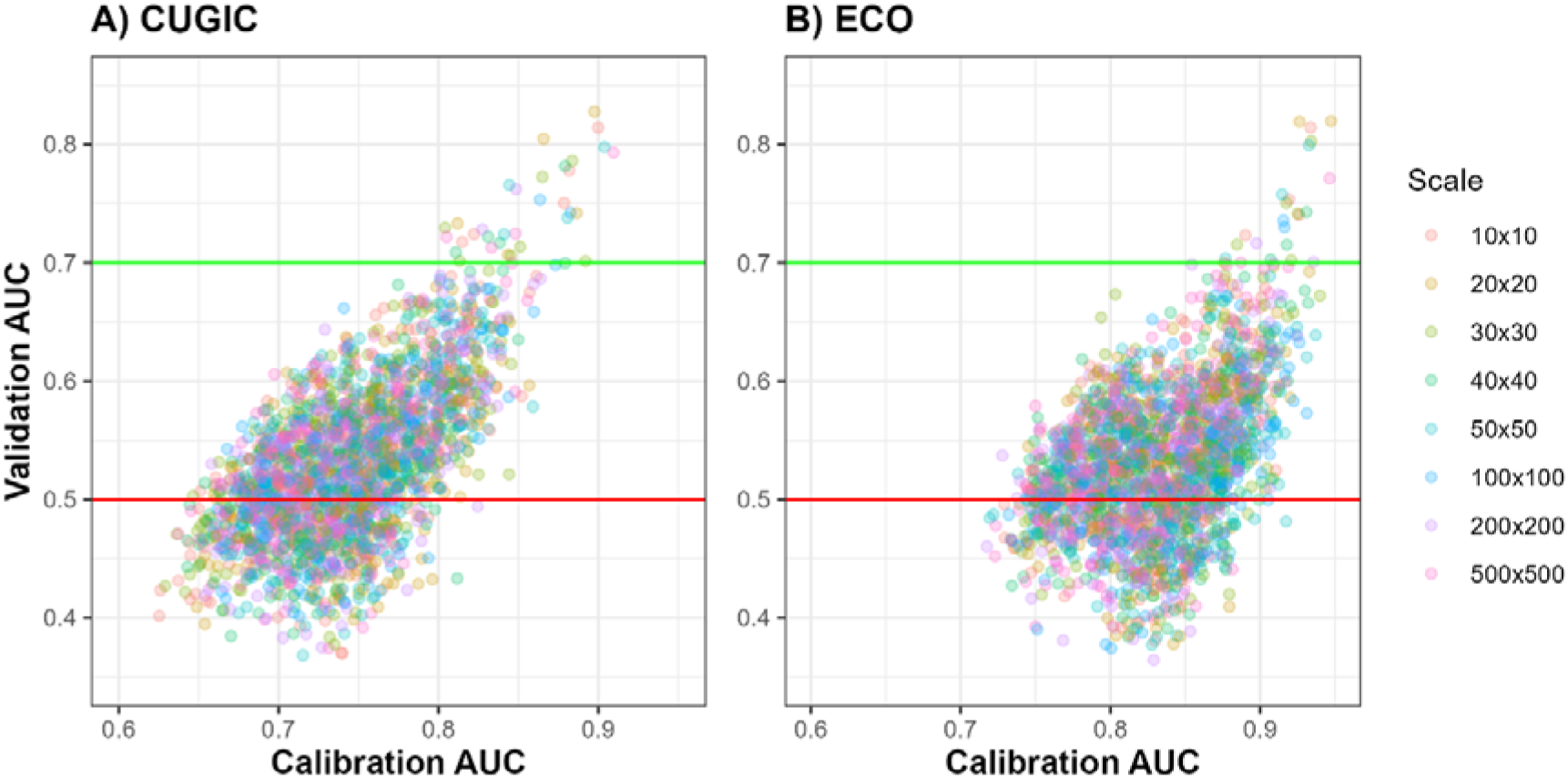
Scatterplot of the AUC values during calibration and validation phase of the species distribution modelling, across eight different spatial scales. Panel A) shows the resulting AUCs from the SDMs with the CUGIC predictor set and panel B) shows the results with the ECO predictor set. The red line shows the performance of 0.50 AUC, which indicates that the SDM has no classification power. The green line shows the performance of 0.70 AUC, where the SDM has a fair or better classification performance.

### 3.3 Global drivers of SDM performance

The linear mixed model assessing the validation AUCs of significant SDMs (n = 2664, 59.3%), shows that dispersal strategy and driver set (CUGIC-model) drive the variation in validation AUC (Table 3). In contrast, the link between scale and AUC is fluctuating between a positive and negative effect, and is rarely statistically significant.

**Table 3.**
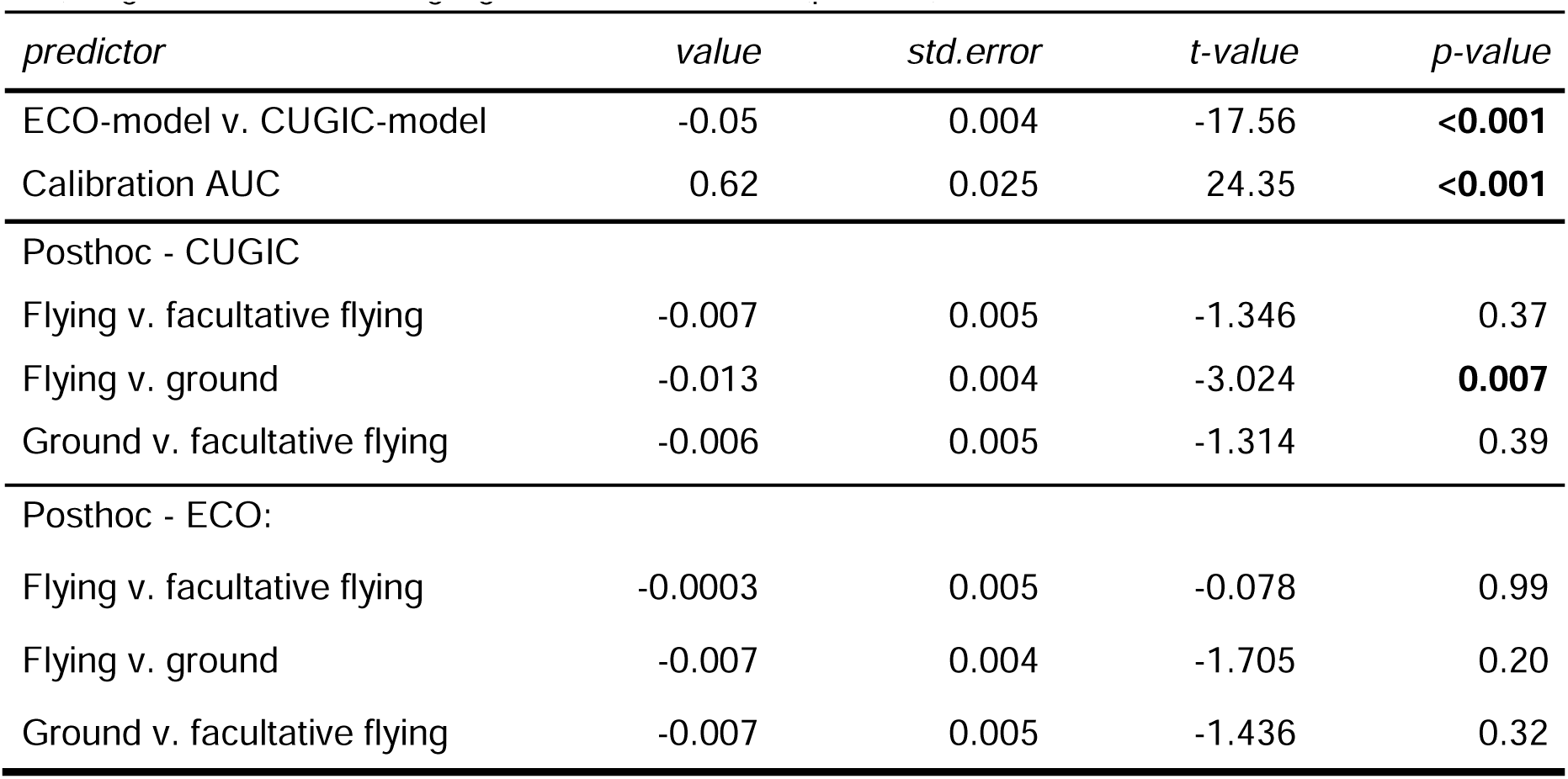
Linear mixed model and the Tukey post-hoc results of AUC by dispersal strategy and driver set. The results of the scale term were omitted due to its inconsistent effect (for full table see S2). Significant terms are highlighted with a bold font (p < 0.05).

| *predictor* | *value* | *std.error* | *t-value* | *p-value* |
| --- | --- | --- | --- | --- |
| ECO-model v. CUGIC-model | -0.05 | 0.004 | -17.56 | **<0.001** |
| Calibration AUC | 0.62 | 0.025 | 24.35 | **<0.001** |
| Posthoc - CUGIC |  |  |  |  |
| Flying v. facultative flying | -0.007 | 0.005 | -1.346 | 0.37 |
| Flying v. ground | -0.013 | 0.004 | -3.024 | **0.007** |
| Ground v. facultative flying | -0.006 | 0.005 | -1.314 | 0.39 |
| Posthoc - ECO: |  |  |  |  |
| Flying v. facultative flying | -0.0003 | 0.005 | -0.078 | 0.99 |
| Flying v. ground | -0.007 | 0.004 | -1.705 | 0.20 |
| Ground v. facultative flying | -0.007 | 0.005 | -1.436 | 0.32 |

In general, the SDMs with the ECO-type showed statistically significantly lower AUC values compared to the CUGIC-type SDMs, which implies the additional variables do not include relevant information for the SDMs (value = -0.05, p < 0.001; Figure 3). However, this effect interacts with the dispersal strategies. The post-hoc analysis shows that for the CUGIC model, flying invertebrates show a consistently lower AUC than ground-dwelling invertebrates (estimate = -0.013, p = 0.007). In contrast, for the ECO-model, we found that none of the dispersal groups performed significantly better than the other. These results suggest that the lower statistical power of the ECO models caused a loss in statistical power to distinguish effects of differences in dispersal capacity.

**Figure 3.**
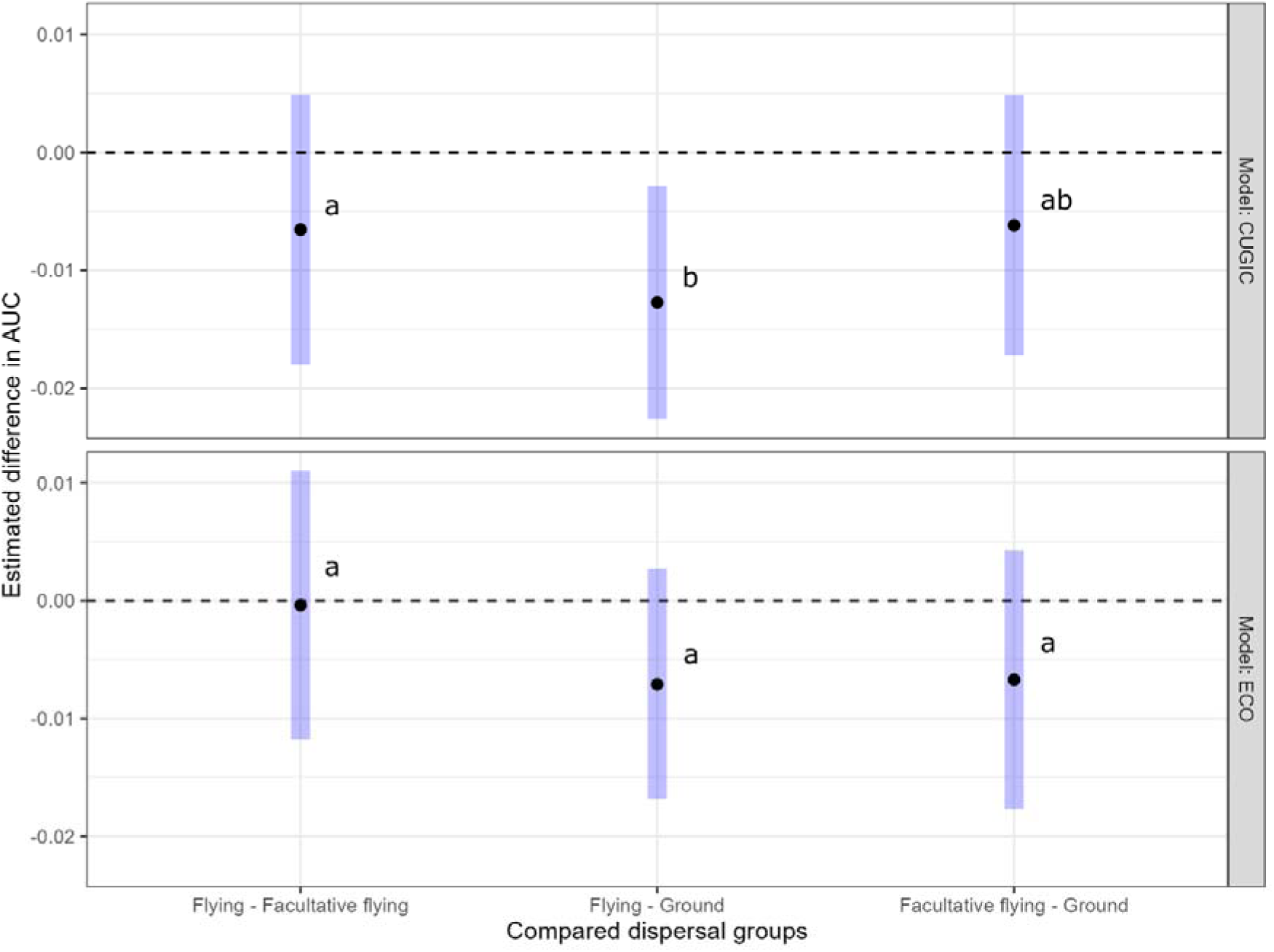
Estimated differences in SDM performance (AUC) across predictor sets (CUGIC/ECO), comparing groups with different dispersal capacities. The horizontal dotted line in both panels indicates no difference between the compared groups. The black dot shows the estimated difference between the groups, and the blue box shows the 95% confidence interval.

### 3.4 Variable importance of the SDMs

The ANOVA and post-hoc analysis on variable importance of significant SDMs (n = 2664, 59.3%), shows that among the CUGIC and ECO models, vegetation density was the most important variable for predicting species presence (Figure 4). In the CUGIC models, vegetation density was the most important variable for ground and facultative flying invertebrates, while flying invertebrates SDMs were equally dependent on LULC. A similar pattern was visible in the ECO models, where vegetation density was the most important variable for ground and facultative flying invertebrates. However, the distance to water was more important to flying invertebrates, followed by vegetation density.

**Figure 4.**
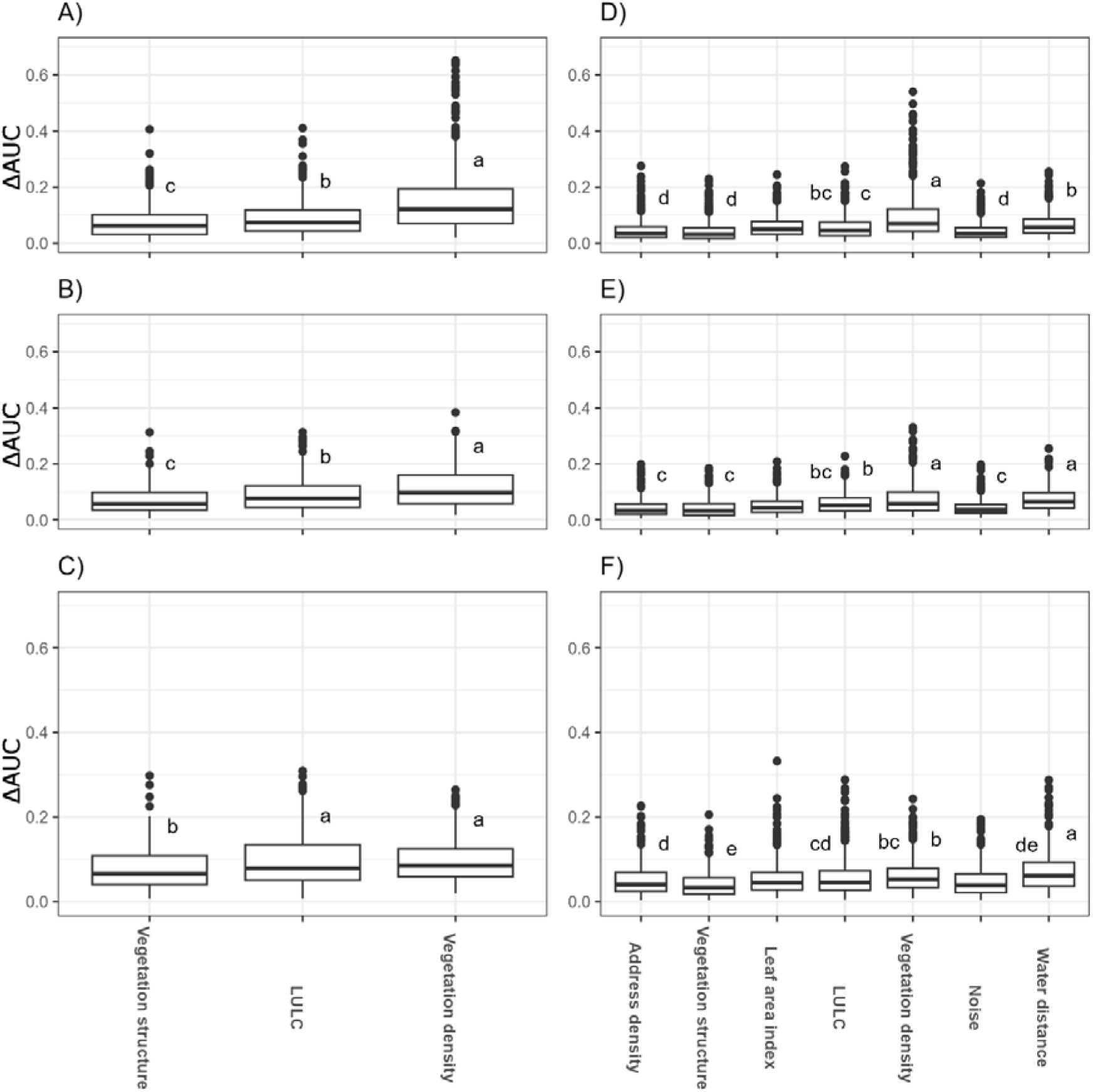
Variable importance scores by model and dispersal group. The left panels, A) — C), show the results for the CUGIC SDMs and right panels, D) — F), the ECO SDMs. The first row, A) and D), show the results for the ground invertebrates. The second row, B) and E), show the results for the facultative flying invertebrates. The third row, C) and F), show the results for the flying invertebrates. The AUC variable importance displays the effect of exclusion of the variable in the model on the performance of the SDM. Letters next to the boxes are corresponding to a post-hoc analysis, grouping variables that are not significantly different from each other (p > 0.05). The alphabetical order indicates which group is the most important for the SDM. LULC = land-use/landcover

In contrast, the least important variables for predicting species presence were both noise and address density. The noise variable scored consistently as least important variable for all the invertebrate SDMs. The address density scored lowest for the ground-dwelling and facultative flying invertebrates, but performed slightly better in the case of flying invertebrates. These results on most and least important variables indicate that ecological indicators are likely important for predicting invertebrate distribution, although different in strength by taxa, while indicators on anthropogenic pressures are unlikely to be important for assessing invertebrate distribution.

### 3.5 Heatmaps of species distributions

The validation threshold of AUC-score > 0.70 resulted in the inclusion of 14 different species containing 57 SDMs explaining their distribution well. From the best performing models per species, we found most SDMs were aggregated to 20m^2^ resolution (n = 5) and were based on the CUGIC driver set (n = 11). The SDMs explaining species distribution well mostly came from the invertebrate bulk metabarcoding data (n = 9). These results indicate that indicators on urban green infrastructure at a local resolution were the best at modelling the observed urban species.

The resulting distribution heatmaps show niches scattered across The Hague (Fig. 5). For example, all OTUs identified best as *Paristoma notabilis* and *Philoscia moscrum* strongly gravitated towards forested regions. Their most likely habitat shows strong overlap in their distribution, indicating that they are similarly affected by changes to the environment. In contrast, both OTUs identified as *Sarcoptiformes* species do not overlap with the other two species, and their distribution is either more present in the urban center or on the outskirts of the city. The *Trixoscelis obscurella* species visually also seem to tend have similar hotspot closer to the city center compared to the ground-dwelling invertebrates. Interestingly, the maps for the *Hymenoptera, Entobryomorpha* and *Mesaphorura* species all shows coarser grids, indicating their best model includes data from a larger scale, even though their dispersal styles are different. These results reflect that urban species occupy different habitats across the city but are also able to sharing identical hotspots. These results indicate species-specific SDMs may provide better overview of several urban invertebrate species their distribution compared to broad biodiversity patterns.

**Figure 5.**
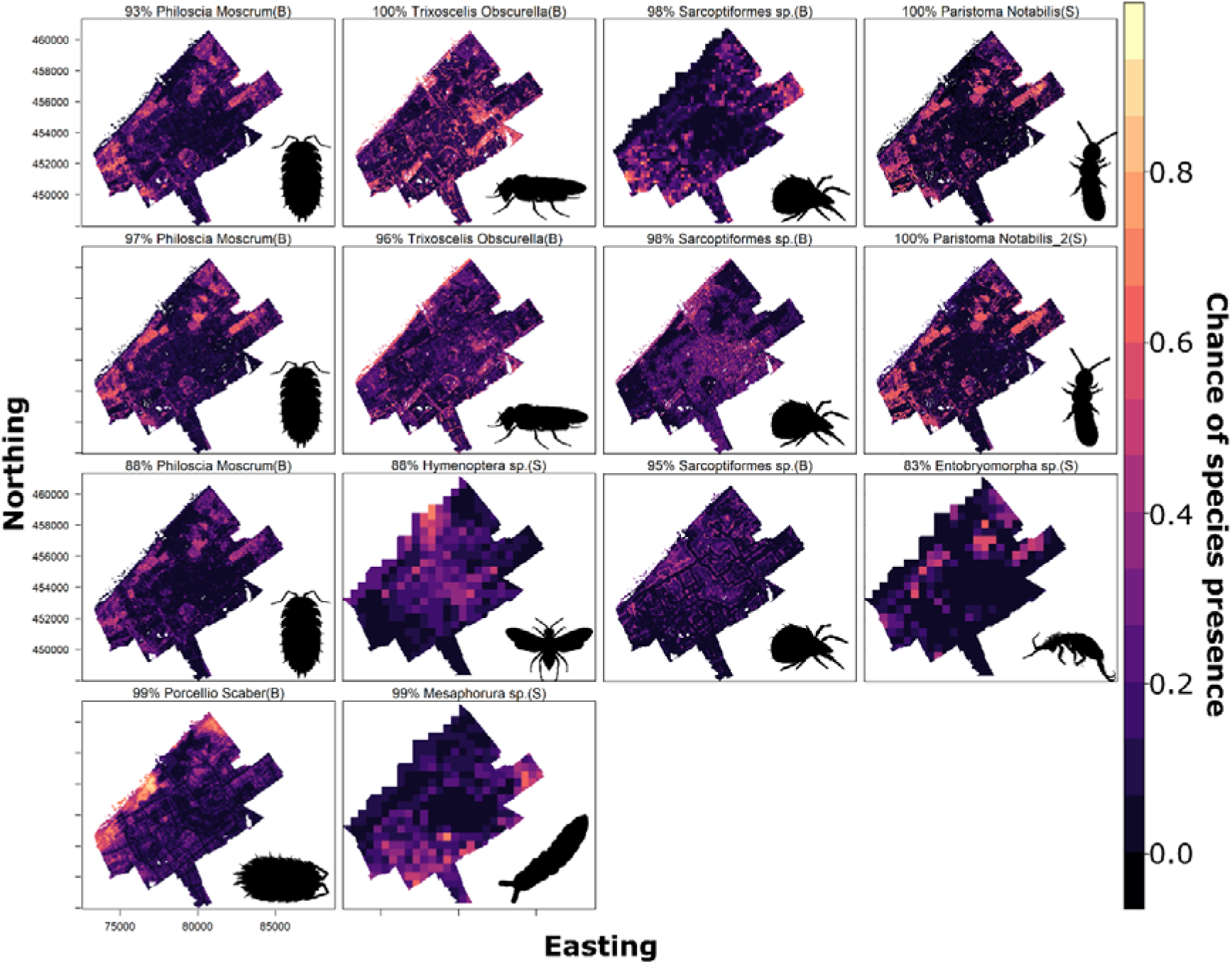
Predicted distributions of the 14 species where the SDMs had an AUC > 0.70. The species names indicate the best match between the modelled OTU and the BOLD reference data, where the percentages indicate the overlap between the OTU and the reference data. An (S) indicates the SDM is based on data from the soil eDNA dataset and a (B) indicates the SDM is based on the data from the invertebrate bulk.

## 4. Discussion

The presented results are the first glimpse at the potential for highly detailed coverage of invertebrate species and for understanding their distributions in urban environments using a combination of DNA-based data and SDMs. Our results show that vegetation and water are most important predictors for the distribution of urban invertebrate species, while indicators on urbanisation such as address density and noise are least important. These results indicate that within the urban environment invertebrate diversity seems to be driven largely by the presence of habitat (water and vegetation), instead of the quality of the habitat(s) (noise and address density). We argue that a combination of DNA-based data and SDMs can play a substantial role in assessing the importance of spatial features and their role in the community assembly of the urban biodiversity.

### 4.1 Ecological implications

The utilization of DNA-based techniques for inferring species presence presents a novel avenue for rapidly assessing biodiversity with extensive coverage of species. Our results show that incorporating both eDNA and bulk metabarcoding enhances the coverage, showcasing their complementary nature and the high potential for uncovering vast quantities of species previously undetectable or at reduce labour (Kirse et al., 2021). These techniques will become even more powerful in the near future, with reference databases covering more species and techniques to retrieve, extract, amplify and sequence DNA will likely become less biased (Cristescu and Herbert, 2018). Using both soil eDNA and invertebrate bulk metabarcoding, a more complete inventory of the full invertebrate community became available.

The SDMs based on the eDNA data showed that within the urban environment, species distribution predictions depended on indicators of vegetation and water, while anthropogenic pressures show minimal impact on species distributions. Several studies also have shown a similar effect, suggesting that available habitat is impacting species distribution more than anthropogenic influences. For example, this has been demonstrated in Singapore, where the presence of urban vegetation and parks was the main determinant of butterfly species presence (Jain et al., 2021). For weaver birds’ movements in Cape Town, increasing urbanization appears inconsequential while the presence of ecologically crucial wetlands plays a decisive role in the bird’s distribution (Calder et al., 2015). These results are also congruence with two studies that performed a meta-analysis on biodiversity within cities, and showed that habitat in the form of vegetation density, abundance and structure are the most important drivers of urban biodiversity (McKinney, 2008; Beninde et al., 2015). These results on green infrastructure all suggest that sufficient terrestrial and aquatic habitat is the limiting factor to enhancing urban biodiversity, and that creating habitat through urban green spaces is the key to increasing biodiversity in cities.

Interestingly, we do not find anthropogenic pressures such as address density and noise to be important for most urban species in our analysis. Anthropogenic pressures have been shown to be useful for estimating species distributions when rural territories are included. Ample prior studies that cover the full rural to urban gradient even suggest that anthropogenic pressures are a pivotal driver of biodiversity (McKinney, 2008; Jain et al., 2021; Beninde et al., 2023). For example, the results of a study examining 1200 terrestrial species using iNaturalist data from the greater Los Angeles, covering the full rural to urban gradient, shows a pronounced association between species distributions and urbanization (Beninde et al., 2023). Similarly, a review on species richness stated that across the rural to urban gradient species richness change, but mostly due to the turnover from locally unique species, to globe-trotting species (Mckinney, 2008). This suggests that the species found in the urban environment are (pre-) adapted to anthropogenic pressures, and can therefore survive well in the urban environment (i.e. urban exploiter; Kark et al., 2007). These (pre-) adapted species would not be affected by anthropogenic pressures which could explain the lack of meaningful contribution of anthropogenic pressures to our models. Indicators on anthropogenic pressures would be most informative in models that cover areas with little to no human impacts, as we expect that species most sensitive to anthropogenic pressures are present there (e.g. Glisson et al., 2017).

Contrary to expectations, while most SDMs were better than random (59.5%), only a few met the traditional AUC standards for SDM model performance of 0.70 (1.4%). The predictors intended for urban ecosystems—such as vegetation density, noise, and distance to water—overfitted the models rather than enhancing their performance. We used these predictors as they are thought to be important for biodiversity in the urban environment and typically used globally available climate data lack relevance at the local urban scale (Beninde et al., 2015; Morpurgo et al., 2023).

There are two possible explanations for why our models failed to meet the performance threshold (AUC > 0.70). Our first explanation revolves around a potential mismatch in the selection of predictors, which are meant to be informative for explaining biodiversity metrics but may lack species-specific niche information. To model species niches accurately, it is important to consider the life-history of the organism, spatial scale (Aguirre-Gutiérrez et al., 2013; Wunderlich et al., 2022), and its environmental drivers when selecting appropriate predictors (Araújo and Guisan, 2006). In our study, there was no difference in performance across eight different spatial scales, supporting the argument that species-specific drivers of niches might have been missing. Here, we suggest some predictors that may improve urban SDMs performances. Recently developed micro-climatic data could be particularly vital for urban SDMs, with natural systems already showing temperature variations of up to 10°C (Suggitt et al., 2011). Several studies have already shown that micro-climate data at a resolution of 25m^2^ varyingly enhances model performance for many species (Lembrechts et al., 2019). We hypothesize that local climate data may be important in cities, as micro-climates are known to be more extreme, by for example the urban heat island effect. Also, indicators specific to pollinators such as floral availability or nesting substrate, and indicators specific to ground-dwelling invertebrates such as soil quality, pollution levels, and organic matter content, have been suggested to enhance predictions of their distributions (Santorufo et al., 2014; Silva et al., 2014). Our results, alongside existing literature, underscores the necessity of exploring other predictors that may be more informative for the spatial distribution of urban species.

A more radical second explanation for the low performance of most SDMs could be that our species are (pre-)adapted to the urban environment, are free to disperse between patches and act as one meta-community. Given that we have likely accounted for most of the important predictors for our target species, and given that the observed species share important ecological traits such as, resources, habitat, mutualists, neutral theory may explain our poorly performing SDMs (Leibold and McPeek, 2006). Interestingly, urban environments are known to biotically homogenise their species on the basis of species being generalists in their ecological needs, easily adapting to novel food sources and habitats, while also having high dispersal capacity (Kark et al., 2007). The congregation of species with similar traits in the urban environment makes a strong case for it being a model system for neutral dynamics. Given neutral theory, the assembly and distribution of urban invertebrate species would be primarily driven by birth, death, and immigration instead of the environment (Bell, 2000). The outcomes of such a neutral system would align with our results, suggesting that habitat availability, i.e. the amount of green space, acts as a limiting factor rather than quality characteristics (such as noise, address density) of urban green infrastructure.

### 4.2 Implications for urban planning

Given that environmental predictors on green infrastructure and water were identified as the most important drivers for urban invertebrate species distributions, investing in the protection and enhancement of these infrastructures is imperative. This is in line with other work from biodiversity studies (Beninde et al., 2015), but also aligns with research on climate adaptation and human health that show that higher amounts of urban nature enhance benefits (Bratman et al., 2019; Remme et al., 2021; Morpurgo et al., 2023). In combination with our finding that anthropogenic pressures such as address density and noise do not substantially affect urban invertebrates distributions, this indicates that plans that could add urban green spaces at a large scale by incorporating highly designed green such as green roofs, street greenery, and rain gardens likely have a strong positive effect on urban biodiversity. Our results therefore strengthen the plea for urban planners and policymakers to prioritize the development and preservation of green spaces, parks, gardens, and other potential habitats within urban areas to support and enhance urban invertebrate biodiversity alongside human-centred benefits.

## 5. Conclusion

Enhancing invertebrate biodiversity is important for safeguarding biodiversity targets and sustaining ecosystem functions essential to humans. For urban invertebrate biodiversity, our analysis shows that environmental predictors on habitat availability, food and water are the most important drivers for urban invertebrate species distributions. In contrast, within the urban environment, species distributions do not seem to be impacted by anthropogenic pressures. We argue that DNA-based SDMs allows for the assessment of urban invertebrate species, their distribution and the driving factors of urban community assembly. Further research should prioritize investigating understanding the mechanisms of urban invertebrate distribution as much of it is left unexplained. We urge policymakers and urban planners to invest in more green and blue infrastructure as they were the most important for invertebrate species’ presence.

## Supporting information

Supplementary_1_data_clhs

Supplementary_2_primers

Supplementary_3_extra_model_results

## 6. Acknowledgements

We would like to thank the following people for their contribution during fieldwork, lab work, or consultation: Jenny Lüdtke, Anne Wilts, Klaas Land, Sofie Rasmussen, Rianne van Duinen, Iris de Wolf, Laura Zantis, Stephanie Cap, Mike Slootweg, Ilmer Benda, Mike Lakerveld, Puck Bramer, Meilin Remijn, Sandhya S Jhingoer, Niels Raes, Yali Si, Kat Stewart, Jan Macher, and Kevin Beentjes.

## Data availability

Data is available at repository XXX (Links to authors repository, removed to keep review blind).

