## Supplementary_1_data_clhs for "Invertebrate species distributions in urban ecosystems are driven by habitat availability, not by anthropogenic drivers"

###### Table with variables used in the Conditional Latin Hypercube Sampling

| **Name** | **Description** |
| --- | --- |
| Leaf Area Index | Numeric variable indicating vegetation. Data is retrieved from the SNAP program from Copernicus, using Sentinel S2A satellite imagery (ESA, n.d.). |
| Temperature feel | Numerical variable describing the modeled temperature felt in a certain cell. Data is retrieved from Klimaateffectatlas (Klimaateffectatlas, 2024) |
| Building height | Height of a building using the Digital Surface Model (DSM) from AHN2 available from Google Earth Engine (AHN, 2024). |
| Ownership | A categorical variable describing the ownership of the GI, being either open or private. This data is gathered using the Kadastrale Percelen dataset from the municipality of The Hague (Kadaster, 2022). |
| Distance to water | Numerical variable describing the distance from a point to the nearest in-land water source. Data on water presence was retrieved from the Basisregistratie Grootschalige Topografie (BGT) and both sea and dry ditches were removed from the dataset before assessing the distance (Kadaster, 2022). |
| Distance to sub-motorway | Numerical variable describing the distance from a point to the nearest sub-motorway (<50km/h). Data was retrieved from Geofabrik (n.d.). |
| Distance to motorway | Numerical variable describing the distance from a point to the nearest motorway (>50km/h). Data was retrieved from Geofabrik (n.d.). |
| Distance to tram track | Numerical variable describing the distance from a point to the nearest tram line. Data was retrieved from Geofabrik (n.d.). |
| Distance to train track | Numerical variable describing the distance from a point to the nearest train track. Data was retrieved from Geofabrik (n.d.). |
| CUGIC vegetation | A categorical variable using scaled NDVI and DSM from AHN2 to assess the vegetation structure. This resulted in 24-categories which describe the vegetation density and the height at which it occurs compared to sea-level. |

***Literature***

- AHN. (2024). Actueel Hoogte Bestand Viewer. Actueel Hoogte Bestand 2. <https://www.ahn.nl/dataroom>
- ESA. (n.d.). Sentinel Application Platform (SNAP). ESA.
- Kadaster. (2022). BAG. In Basisregistratie Adressen en Gebouwen v2.
- Klimaateffectatlas, 2024. Kaartviewer. <https://www.klimaateffectatlas.nl/nl/kaartviewer>
- Jochen, T., Frederik, R., Christine, K., Amanda, M., Philip, B., & Michael, R. (n.d.). Geofabrik. (n.d.). Geofabrik - Free Geodata.
