## Supplementary_2_primers for "Invertebrate species distributions in urban ecosystems are driven by habitat availability, not by anthropogenic drivers"

**Supplementary information on primers**. Shown are the primers used and the exact sequences of basepairs.

| **Primer name** | **Sequence 5’ 🡪 3’ orientation** |
| --- | --- |
| BF1 | ACACTCTTTCCCTACACGACGCTCTTCCGATCTACWGGWTGRACWGTNTAYCC |
| BR2 | GTGACTGGAGTTCAGACGTGTGCTCTTCCGATCTTCDGGRTGNCCRAARAAYCA |
| mlCOIintF | ACACTCTTTCCCTACACGACGCTCTTCCGATCTGGWACWGGWTGAACWGTWTAYCCYCC |
| dgHCO2198 | GTGACTGGAGTTCAGACGTGTGCTCTTCCGATCTTAAACTTCAGGGTGACCAAARAAYCA |
