## Supplementary_3_extra_model_results for "Invertebrate species distributions in urban ecosystems are driven by habitat availability, not by anthropogenic drivers"

###### Table with results from the linear mixed model predicting AUC by scale, model type and dispersal group. Estimates are the difference in AUC between scales of 10x10m and the scale of the term. The dispersal groups compared are flight (as basis for comparison) against facultative flying and ground. The X between a dispersal group and scale indicate a modelled interaction. In bold are the statistically significant terms at p < 0.05.

| Term | Estimate | Std.error | p-value |
| --- | --- | --- | --- |
| scale 20 | 0.003 | 0.003 | 0.38 |
| scale 30 | -0.004 | 0.004 | 0.32 |
| scale 40 | -0.003 | 0.004 | 0.38 |
| scale 50 | 0.00006 | 0.004 | 0.98 |
| scale 100 | -0.006 | 0.004 | 0.08 |
| scale 200 | -0.001 | 0.004 | 0.72 |
| scale 500 | 0.004 | 0.004 | 0.21 |
| facultative flying X scale 20 | -0.009 | 0.006 | 0.12 |
| facultative flying X scale 30 | -0.005 | 0.006 | 0.34 |
| facultative flying X scale 40 | -0.013 | 0.006 | **0.03** |
| facultative flying X scale 50 | -0.010 | 0.006 | 0.10 |
| facultative flying X scale 100 | -0.004 | 0.006 | 0.49 |
| facultative flying X scale 200 | -0.006 | 0.006 | 0.33 |
| facultative flying X scale 500 | -0.012 | 0.006 | 0.06 |
| ground X scale 20 | -0.009 | 0.005 | 0.08 |
| ground X scale 30 | 0.006 | 0.005 | 0.20 |
| ground X scale 40 | -0.005 | 0.005 | 0.37 |
| ground X scale 50 | -0.008 | 0.005 | 0.12 |
| ground X scale 100 | -0.008 | 0.005 | 0.11 |
| ground X scale 200 | -0.005 | 0.005 | 0.30 |
| ground X scale 500 | -0.0006 | 0.005 | 0.90 |
